# Network analysis to evaluate the impact of research funding on research community consolidation

**DOI:** 10.1101/534495

**Authors:** Daniel J. Hicks, David Coil, Carl G. Stahmer, Jonathan A. Eisen

## Abstract

In 2004, the Alfred P. Sloan Foundation launched a new program focused on incubating a new field, “Microbiology of the Built Environment” (MoBE). By the end of 2017, the program had supported the publication of hundreds of scholarly works, but it was unclear to what extent it had stimulated the development of a new research community. We identified 307 works funded by the MoBE program, as well as a comparison set of 698 authors who published in the same journals during the same period of time but were not part of the Sloan Foundation-funded collaboration. Our analysis of collaboration networks for both groups of authors suggests that the Sloan Foundation’s program resulted in a more consolidated community of researchers, specifically in terms of number of components, diameter, density, and transitivity of the coauthor networks. In addition to highlighting the success of this particular program, our method could be applied to other fields to examine the impact of funding programs and other large-scale initiatives on the formation of research communities.

## Introduction

In 2004, the Alfred P. Sloan Foundation launched a program focusing on the “Microbiology of the Built Environment”, sometimes known as “MoBE”. The aims of this program were to catalyze research on microbes and microbial communities in human built environments, such as homes, vehicles, and water systems; and to develop the topic into a whole field of inquiry. Prior to 2004, many new developments (e.g., major advances in DNA sequencing technology) had catalyzed innovation in studies of microbes found in other environments (e.g., those living in and on humans and other animals, those found in the soil, those found in the oceans), but these innovations had not spread rapidly enough to studies of the microbes in the built environment. Similarly, many developments had occurred in studies of the built environment (e.g., the spread of low cost sensor systems), but focus had not yet been placed on the living, microbial components of built environments. This is not to say there had been no studies on the MoBE topic prior to 2004, but rather that the pace of advances in the area were modest at best compared to advances in other areas of microbiology and built environment studies. The MoBE area was founded on the belief that institutionally supported, integrated, trans-disciplanary scientific inquiry could address these shortfalls and lead to major benefits in areas such as indoor health, disease transmission, biodefense, forensics, and energy efficiency.

The Sloan Foundation’s program ultimately lasted 15 years and invested more than $50 million on work in the MoBE field. A key goal of this program was to bring together the highly disparate fields of microbiology (especially the area focused on studies of entire ecosystems of microbes) and building science (e.g. with a focus on building, maintaining, regulating, and studying built environments) with their different approaches, cultures, incentives, and rewards. Grants were given to many projects and a diverse collection of people covering many fields including microbiology, architecture, building science, software development, and meeting organization (a list of all grants from the program can be found at https://sloan.org/grants-database?setsubprogram=2). The products of these grants included a diverse collection of programs and projects, dozens of new collaborations, many novel and sometimes large data sets on various MoBE topics, new software and tools for MoBE studies, and hundreds of scholarly publications.

Recent reviews of the state of the field (e.g. [1][2]) have qualitatively highlighted the success of this program. In this paper we report a quantitative assessment of the Sloan MoBE program and the MoBE field using a network analysis of scholarly literature. Specifically, the aim of this study was to compare the community of researchers funded by the Sloan Foundation’s MoBE program to their scientific peers. If the Sloan Foundation’s program was successful at cultivating a new research community around MoBE topics, we hypothesized that we would see the evolution of an increasingly dense and more tightly connected network over the duration of the funding program.

## Methods and Materials

### Corpus Selection

Publications funded by the Sloan Foundation’s MoBE program provided the starting point for our data collection and analysis. We evaluate the effect of this program by analyzing these publications in the context of previous work by the same authors, as well as a “control” or comparison set of authors working in the same general areas. We identify the comparison set as authors publishing frequently in the same journals as MoBE-funded publications.

### Identifying Sloan Foundation-Funded Publications

A list of publications associated with the Sloan Fundation-funded MoBE program was compiled through a combination of strategies. An initial set of papers was identified by manually searching for acknowledgement of Sloan Foundation funding in any publications authored by the grantees during the program period. Additional publications were identified by searching Google Scholar for relevant MoBE papers and identifying those authored by grantees during the program period. Finally, each grantee (as well as sometimes their lab members (n=∼50)) was contacted directly and asked whether the publication list we had for them was both accurate and complete. This feedback led to some publications being removed from the list (as having not derived from the Sloan Foundation’s program) and others being added. In addition, we posted requests for feedback in various social media settings (e.g., blogs, Twitter) asking for feedback on the list. The final list contained 327 publications. 20 of these publications did not have digital object identifiers (DOIs) on record and were excluded from further analysis.

### Identifying Peer Authors

We sought to compare MoBE researchers to peers who were not funded by the MoBE program, in order to control for ordinary developments in both individual careers (e.g., more senior researchers are likely to have more collaborators) and research communities (e.g., more researchers are trained and join the community). In what follows, researchers funded by the Sloan Foundation’s program are referred to as the “collaboration” authors; their peers are the “comparison” authors.

Several methods were considered for developing this comparison set. Keyword searches were judged to be too noisy, producing significant numbers of false positive and false negative matches, as well as highly sensitive to the particular keywords used. Forward-and-backward citation searches using the 307 MoBE articles (compare [3]) produced lists on the order of 1,000,000 publications, which was judged to be impractically large. As an alternative, peer authors were identified as authors who are highly prolific in the same journals as the 307 MoBE articles.

Specifically, using the rcrossref package [4] to access the Crossref API (application programming interface; https://github.com/CrossRef/rest-api-doc), metadata were retrieved for 572,362 articles published in 111 journals between 2008 and 2018 inclusive. (*PLOS One* was dropped prior retrieving these metadata, due to its general nature and extremely high publication volume.) 14 journals published at least 10,000 articles during this time period; these appeared to be high-volume, general or broad-scope journals, such as *Science* or *Environmental Science & Technology*. The 345,546 articles from these 14 journals were removed, leaving 226,816 articles from 97 journals. Because Crossref does not provide any standardized author identifiers, simple name matching was used to estimate the number of articles published by each author. (This method means “Maria Rodriguez” and “M. Rodriguez” would be counted as different authors at this stage.) The same method was used to roughly identify authors of MoBE-funded papers. After filtering out authors of MoBE-funded papers, the 1,000 most prolific authors were selected as candidates for the comparison set. See fig. 1.

**Figure 1:**
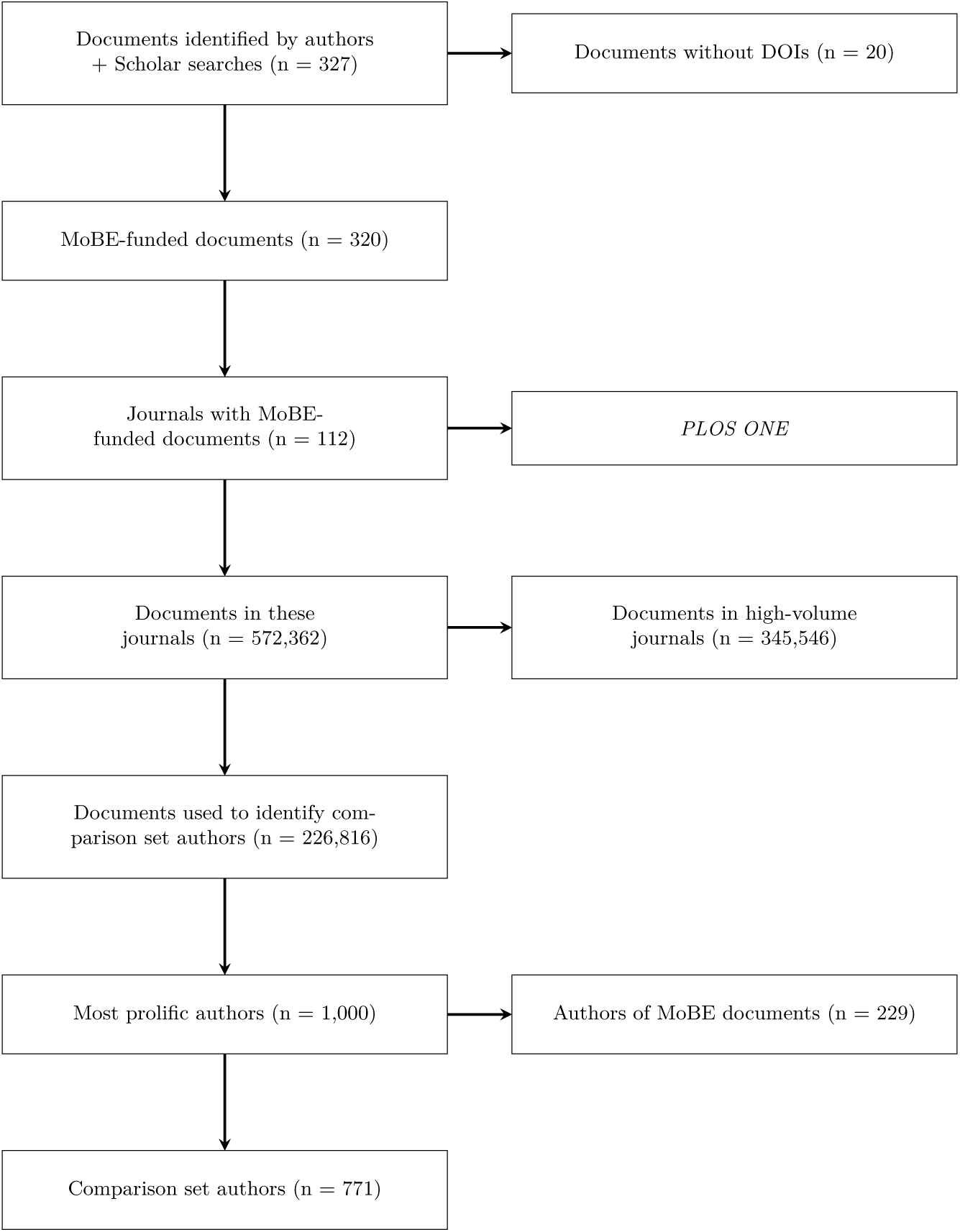
Flow diagram for comparison set construction.

Next, to retrieve standardized author identifiers, a covering set of papers was identified; each candidate appeared as an author of at least one paper in the covering set. Metadata for these papers was retrieved from the Scopus API (https://dev.elsevier.com), which incorporates an automated author matching system and standardized identifiers, referred to as author IDs. These author IDs were used to characterize researchers as members of the MoBE collaboration or comparison set. Collaboration authors were defined as any author who either (a) was an author of at least two MoBE-funded papers or (b) was the author of at least one MoBE-funded paper and appeared in the candidates list (total n=393 distinct names for the collaboration; 438 distinct author IDs). Candidates for the comparison set were removed if they were classified as part of the collaboration (total n=771 distinct author IDs for the comparison set). (In what follows, we do not distinguish between authors and author IDs.)

### Author Histories and Collaboration Network Analysis

Author histories (up to 200 more recent publications since 2003 inclusive) for all 1,209 authors were retrieved using the Scopus API. These histories include both MoBE-funded and non-MoBE-funded papers, published in all journals indexed by Scopus. This resulted in an analysis dataset of 81,312 papers. Besides standard metadata, each paper was identified as MoBE-funded (or not).

The resulting dataset of 81,312 papers was used as the basis for constructing time-indexed collaboration networks. Each author forms a node (distinguished by author ID); edges correspond to papers published in a given year, so that two authors are connected by an edge for a given year if they coauthored at least one paper published in that year. All collaboration authors had at least one edge; 73 comparison authors did not have at least one edge, and were dropped from the network analysis (remaining comparison n = 698). Authors who collaborated on multiple papers in a given year were connected with multiple edges, except when calculating density (see below).

After constructing the combined (collaboration + comparison) network, separate cumulative-annual networks were constructed for each set of authors. For example, two authors would be connected in the 2011 network if and only if (1) they were in the same author set and (2) they had coauthored at least one publication between 2003 and 2011 inclusive. Cumulative networks were used to reduce noise in the most recent years, due to incomplete data for 2018 and as the Sloan Foundation’s funding program was starting to wind down. Analyzing separate cumulative networks allows the examination of the development of research communities through time and between the author sets.

[5] offer an approach to identifying the consolidation of research communities using topological statistics of networks. Specifically, they propose that community formation can be measured in terms of giant component coverage and mean distance or shortest path length: increasing coverage combined with decreased distance indicates community consolidation. We extended this approach by calculating a total of eight network topological statistics and directly comparing the two author sets. Specifically, we calculated the number of authors, number of components, coverage of the giant component (as a fraction of authors included in the largest component), entropy (*H*) of the component size distribution, diameter, density (fraction of all possible edges actually realized), mean distance, and transitivity in each year.

Number of authors simply measures the total size of each network. Because these are cumulative networks, the number of authors necessarily increases. The number of components, coverage of the giant component, and entropy of the component distribution are measures of the large-scale structure of the network. More components indicate that the network is divided into subcommunities that do not interact (at least in terms of coauthoring papers); fewer components indicates consolidation of the research community. Giant component coverage and entropy measure the relative sizes of these different components; higher giant component coverage and lower entropy indicate that more authors can be found in a single component, which in turn indicates research community consolidation.

Diameter, density, and mean distance can be interpreted as measures of the ability of information to flow through the network. Lower diameter, higher density, and lower mean distance indicate that it is easier for information to move between any two given researchers, as there are fewer intermediary coauthors and a higher probability of a direct connection. These therefore indicate research community consolidation.

Transitivity is an aggregate measure of the local-scale structure of the network. Low transitivity indicates that the network is comprised of loosely connected clusters; there is collaboration across groups of researchers, but it is relatively rare. High transitivity, by contrast, indicates that the network cannot be divided into distinguishable clusters. High transitivity therefore indicates research community consolidation.

To analyze the robustness of these observed statistics against data errors or missingness, perturbed networks were generated for each year by randomly switching the endpoints of 5% of edges.

All data collection and analysis was carried out in R [6]. Complete data collection and analysis, as well as the list of MoBE-funded publications, is available at https://doi.org/10.5281/zenodo.2548840.

## Results/Discussion

### Qualitative Analysis

The development of the combined network is shown in Fig. 2. MoBE-funded authors and papers are shown in blue; non-MoBE-funded authors and papers are shown in red. All together, we believe that Fig. 2 shows the consolidation of the MoBE collaboration within a consolidating larger research community.

**Figure 2:**
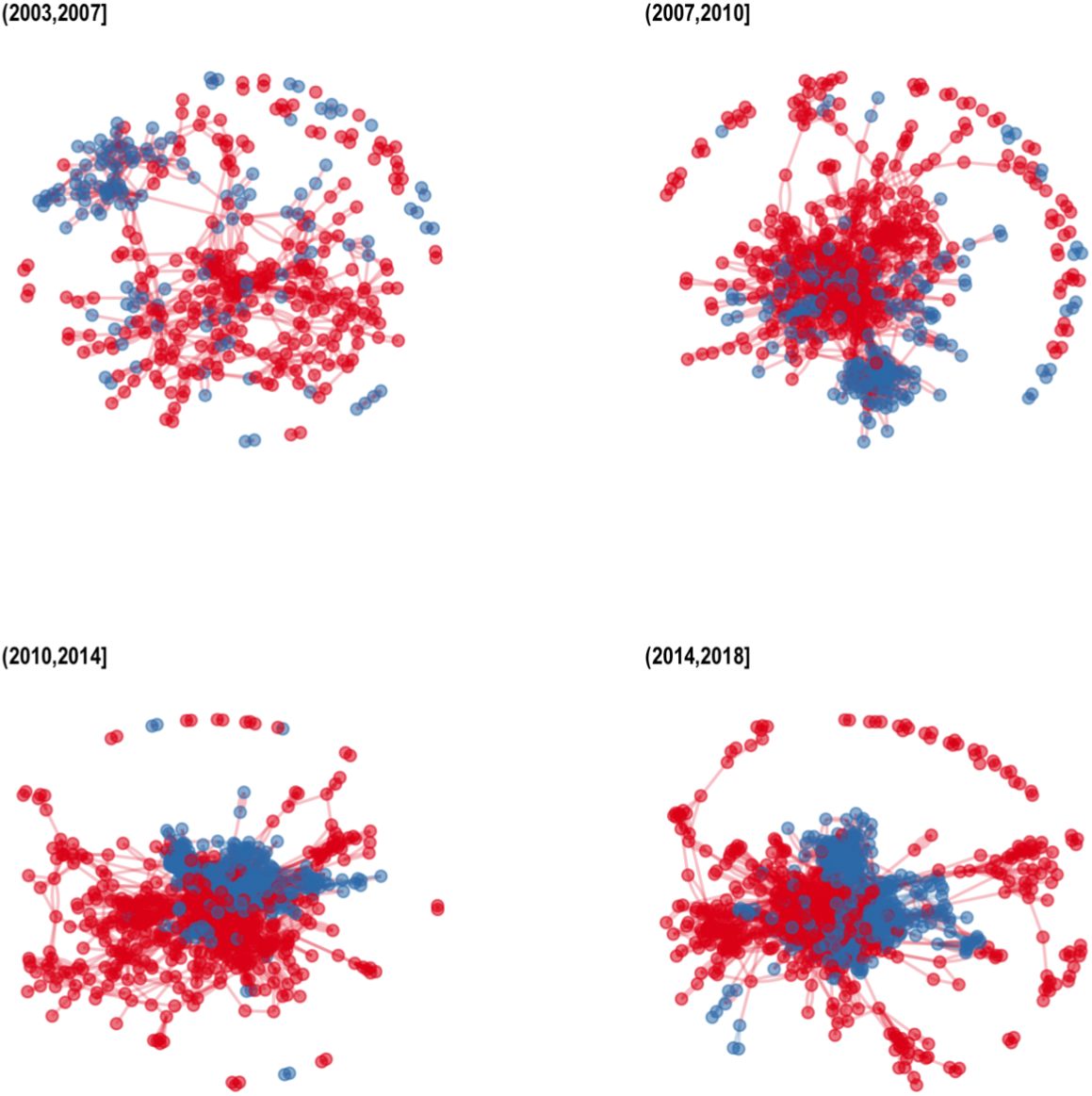
Consolidation of the MoBE collaboration over time. Panels show time slices (non-cumulative) of the giant component of the combined coauthor network. Blue nodes and edges are MoBE authors and papers; red nodes and edges are non-MoBE authors and papers. Network layouts are calculated separate for each slice using the Fruchterman-Reingold algorithm with default values in the igraph package.

Prior to the beginning of the MoBE funding in 2008, a subset of MoBE researchers are actively working with each other; but many MoBE researchers are isolated in this network. Qualitatively, the combined network has a sparse “lace” structure, with many long loops, as well as an “archipelago” of numerous small disconnected components.

During the early years of the funding period (2008-2010), a tighter cluster of MoBE researchers appears on the margins of the overall research community; but many MoBE researchers can be found scattered among the comparison authors and in disconnected components. The combined network has a “hairy ball” appearance, with a dense central “ball” and many peripheral “hairs,” and again an extensive “archipelago.”

In 2011-2014, the combined network appears as a dense “blob” or “clump” with little qualitative structure. The MoBE collaboration appears as a coherent “sub-blob,” with very few MoBE researchers found outside this “sub-blob.” We infer that this indicates that the MoBE collaboration is highly integrated within the larger community.

The combined network is qualitatively similar in 2015-2018. This network may have sightly more internal structure than the 2011-2014 network, suggesting the possibility that the combined research community may be splitting into subcommunities. However, because qualitative features of a visualized network are heavily dependent on the visualization method, this qualitative difference should not be overinterpreted. Data on the development of the community over the next few years would be necessary to examine rigorously the possibility of splitting.

Note that a few comparison set authors remain in small disconnected components even in the final time slice. These likely reflect “false positives” in the construction of the comparison set: authors who appear relatively frequently in the same journals as the MoBE publications, but do not actually conduct research in relevant research areas. We manually identified some such false positives, including authors of news stories in journals such as *Current Biology* or *Nature Biotechnology* as well as a few neuroscientists.

### Quantitative Analysis

Fig. 3 shows statistics over time for the cumulative collaboration networks in each comparison set. Overall, both the MoBE research community and the comparison research community consolidated over time; but the MoBE research community consolidated faster and more thoroughly than the comparison set.

**Figure 3:**
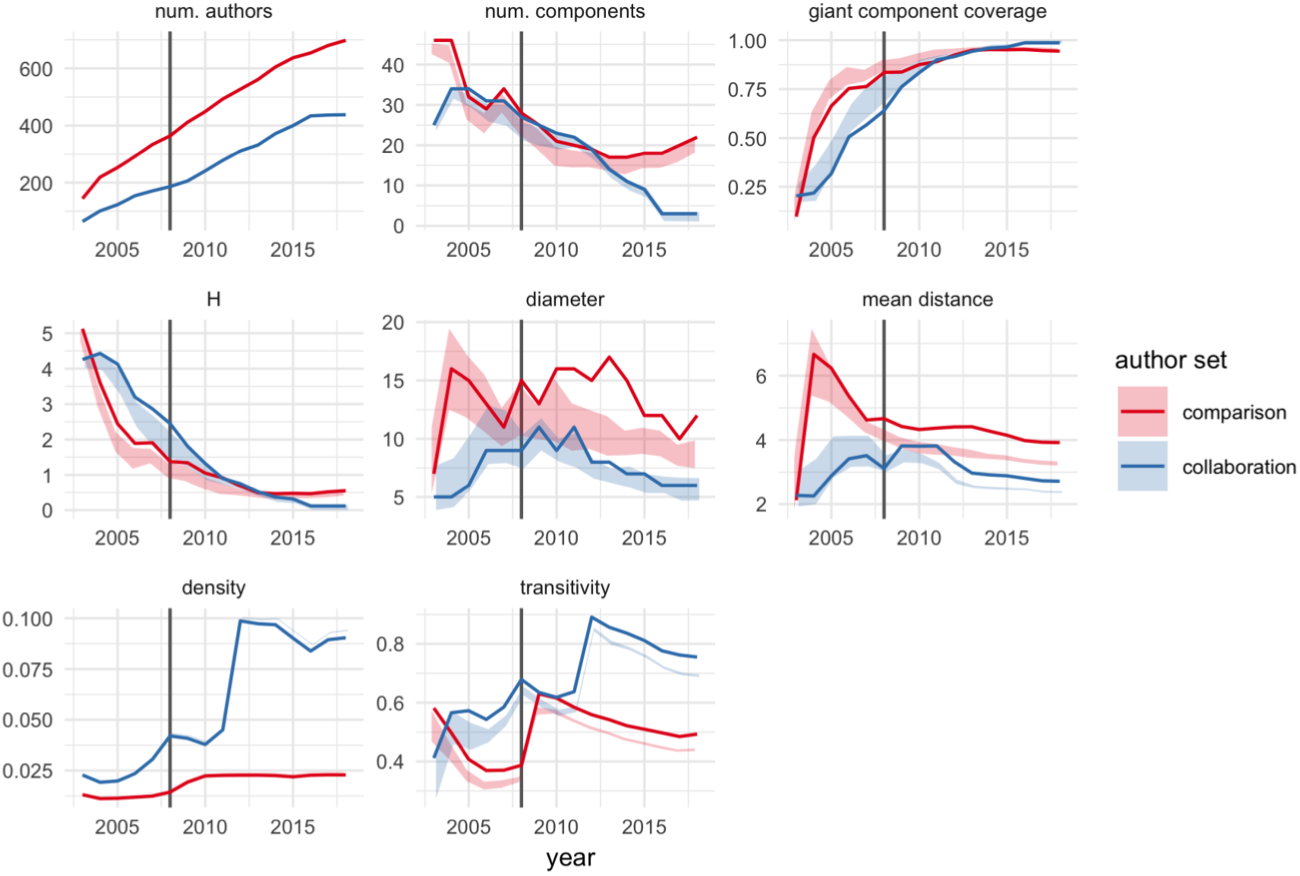
Network statistics over time. See text for explanation of statistics. Solid lines correspond to observed values; shaded ribbons correspond to 90% confidence intervals on perturbed networks, where 5% of the observed edges are randomly rewired while maintaining each node’s degree distributions. 100 rewired networks are generated for each author set-year combination. Blue corresponds to the MoBE collaboration; red corresponds to the peer comparison set of authors. Solid vertical line indicates 2008, the year MoBE funding began.

The most notable differences between the two author sets appear with the number of components, diameter, density, and transitivity. The comparison set stabilizes at 15-20 distinct components, while the MoBE collaboration approaches just two components. However, for both author sets giant component coverage approaches 1 and *H* approaches 0, indicating that both networks contain a single giant component; the comparison set simply has several disconnected components with isolated researchers. As observed in the qualitative analysis, we believe this is plausibly due to “false negatives” in constructing the comparison set. The remaining statistics are generally robust to the inclusion of such “false negatives.”

Diameter remains between roughly 10 and 17 for the comparison set, with a notable decrease after 2013. The MoBE collaboration diameter is consistently less, and in particular is below 10 after 2011.

However, diameter might be criticized as sensitive to network size. The relatively low diameter of the MoBE collaboration might be explained by the fact that this network has about half as many researchers as the comparison set.

By contrast, density and transitivity are normalized against network size. Transitivity and density peak near 90% in 2012, indicating that at this time the connected components of the MoBE collaboration have almost no internal structure: everyone involved in the collaboration in 2012 is working directly with almost everyone else. Transitivity and density then drop somewhat, but still remain remarkably high, indicating a highly interconnected research community even as the number of authors approaches its peak of just over 400. Transitivity is greater than 60% for both author sets in 2008-2009, but then diverges, dropping to around 50% in the comparison set by 2018.

Shaded regions in Figure 2 indicate that most comparisons between the MoBE and comparison sets are robust to data errors. Diameter and mean distance are somewhat more sensitive than the other statistics; but even here the comparison set statistics are consistently greater than the MoBE set statistics, indicating less consolidation in the comparison set.

## Conclusions

Overall, we believe our results support the hypothesis that the Sloan Foundation-funded researchers consolidated as a community over the course of the program. Whereas at the start of the program there were relatively few connections between researchers, especially across domains, by the end of our study period the network was dense and highly interconnected. In particular, while the Sloan Foundation-funded community was initially less connected than the control community it reached a similar level of consolidation by the end of the study period. This suggests to us that the program was successful in the stated goal of increasing collaboration between researchers, particulary from disparate fields.

We note that the most dramatic differences between the MoBE collaboration and the comparison set could not have been detected using the two statistics calculated by [5], namely, giant component coverage and mean distance. Giant component coverage approached unity for both networks, and the difference in mean distance was relatively small. Mean distance could also be criticized as too sensitive to network size. By contrast, the most striking differences in this case appeared in density and transitivity.

## Acknowledgements

The authors would like to thank the many program grantees who assisted us in refining the list of publications. Also thanks to Julia Maritz for compiling the initial list of publications. Funding for DAC and JAE came from the Alfred P. Sloan Foundation. DJH’s postdoctoral fellowship was funded by a gift to UC Davis from Elsevier; Elsevier played no role in designing, conducting, or interpreting the results of this study.

